# TangentXY: Improved copy-number estimation of sex chromosomes reveals features associated with aneuploidies in chromosomes X and Y across cancers

**DOI:** 10.64898/2026.10.01.755972

**Authors:** Kei Enomoto, Shahab Sarmashghi, Sean Misek, Siyun Lee, Greg Raskind, Stephanie Hoyt, Galen Gao, Davy Deng, Xiaomeng Zhang, Jérémie Kalfon, Gordon Saksena, Andrew Cherniack, Rameen Beroukhim

**Affiliations:** Department of Cancer Biology, Dana-Farber Cancer Institute, Boston, MA, USA; Department of Medical Oncology, Dana-Farber Cancer Institute, Boston, MA, USA; Cancer Program, Broad Institute of MIT and Harvard, Cambridge, Boston, MA, USA; Department of Biomedical Informatics, Harvard Medical School, Boston, MA, USA; Harvard-MIT Health Sciences and Technology Program, Boston, MA, USA; Department of Medicine, Harvard Medical School, Boston, MA, USA; Department of Neurology, Brigham and Women’s Hospital, Boston, MA, USA

## Abstract

Signal-to-noise ratios in copy-number estimates across genomic loci improve with the use of large panels of germline reference data. However, in the case of sex chromosomes, germline samples differ between males and females (males have one X chromosome (chrX) and one Y chromosome (chrY), while females have two chrXs), limiting the size of the reference sets to those that match the sex of the index sample. Frequent mosaic loss of chrY in normal male tissue also leads to biases when not accounted for. Here we report TangentXY, a copy-number normalization method that uses both male and female reference samples, correcting for sex differences in chrX and accounting for chrY mosaicism, to assess sex chromosome copy numbers for samples of either sex. We show that TangentXY maintains sex-specific differences in chrX and chrY while minimizing noise. We also used TangentXY to analyze associations between copy-number alterations in sex chromosomes and cancer characteristics, and found that losses in the chrX but not chrY are correlated with TP53 mutation, that gain of chrX can predict patient prognosis in certain tumor types, and that the vast majority of gains of chrX in males remain active, whereas gains of chrX in females are divided approximately equally between active and inactive chrXs. We also found that loss of chrY correlates with inactivating driver mutations in escape genes in chrX, suggesting a mechanism by which its frequent loss promotes cancers in males.

## Introduction

Despite the importance of somatic copy-number alterations (SCNAs) in sex chromosomes in cancer(Dong *et al.*, 2020; Rubin *et al.*, 2020; Kim *et al.*, 2018; Rubin, 2022), analyses of SCNAs in X chromosome (chrX) and Y chromosome (chrY) have been neglected or intentionally excluded from many cancer studies. SCNAs are typically detected after normalizing cancer sequencing or microarray data to germline genomes, but the natural variation between germline dosages of chrX and chrY between males and females complicates this normalization approach.

One solution is to normalize cancer genomic data only to sex-matched germline genomic data. However, excluding germline genomic data from half the population can harm signal-to-noise ratios. In addition, mosaic copy-number changes of chrY in normal tissues increase in prevalence as men age(Jacobs *et al.*, 1961, 1963; Pierre and Hoagland, 1972), which can impair the use of normal male samples as germline controls. To get the most out of all available normal samples we developed a new approach, “TangentXY”. TangentXY extends upon the existing “Tangent” method for copy-number normalization, which makes use of variations in noise patterns across germline genomic data to accurately control for noise patterns in each cancer sample(Gao *et al.*, 2022), but pre-processes the normal samples to remove sex differences in chrX and account for mosaic loss of chrY. This approach properly normalized the sex chromosome signals and enabled us to evaluate associations between copy-numbers in chrX and chrY and cancer characteristics.

## Materials and Methods

### Arm level copy-number analysis on segmentation data in TCGA

Segmentation files generated from Affymetrix Genome-Wide Human SNP Array 6. 0 with GRCh38 coordinates were downloaded from the NIH GDC Data Portal. The average relative copy-number per chromosome arm was determined from the average signal intensity and segment length in each segment of each sample.

### Generation of raw coverage data

GATK DepthOfCoverage tool was used to generate raw coverage data on TCGA WES bam files (mapped to GRCh37) and CCLE WES bam files (mapped to GRCh38) that are stored in Firecloud/Terra platform (https://firecloud.terra.bio/). DepthOfCoverage outputs values for a set of genomic loci (‘intervals’) representing the hybrid capture targets. The interval file (whole_exome_agilent_1. 1_refseq_plus_3_boosters. targetIntervals. bed) is available in The Cancer Genomics Hub (CGHub) bitbucket (https://bitbucket.org/cghub/cghub-capture-kit-info/src/master/). Detailed methods are described in the previous Tangent paper(Gao *et al.*, 2022).

Raw coverage data were successfully generated for 20,765 samples across 33 tumor types (10,650 normal samples and 10,115 tumor samples) from the TCGA data set and for 326 samples from CCLE (25 non-cancerous fibroblast cell lines and 301 cancer cell lines).

### Pre-processing of raw coverage data for downstream analysis (TCGA data)

Samples and genomic loci (also referred to as “intervals” or “probes”) considered to be of low data quality were removed from the raw coverage data. Samples with zero coverage in more than 5% of 182,568 probes designed for autosomal chromosomes were excluded, leaving 20,436 samples (10,487 normals and 9,949 tumors). Likewise, we excluded probes in autosomes and X chromosomes that had zero coverage in 40% or more of all samples; for Y chromosomes, those that did so in 40% or more of all male samples were excluded. A total of 189,320 probes remained.

Quality checks of normal samples using PCA and other methods showed that there was variation in the signal profile between the normal samples (data not shown), but it was expected that including noise from as many different sources as possible (e. g. different labs, different batches of reagents, different types of samples, etc.) in the normal reference would improve TangentXY performance, so no filtering of the normal samples was performed except for anomalous samples described below.

We excluded individuals who were likely to be mislabeled for biological sex, based on the ratio of chrX to chrY signal intensities in their normal DNA. We also excluded samples that were excluded for the same reasons in similar studies conducted by Sadagopan *et al.* and Qi *et al.*, based on their respective criteria(Sadagopan *et al.*, 2022; Qi *et al.*, 2023). Samples that were determined to have Klinefelter syndrome in Sadagopan’s article were also excluded. This resulted in the exclusion of 75 samples taken from 37 individuals. We also removed samples of unknown gender in the TCGA sample information file, resulting in a final sample size of 10,441(5,317 females and 5,124 males) normal samples and 9,854 (5,055 females and 4,799 males) tumor samples for downstream analyses.

We then log2 transformed the coverage data, scaled by the median autosomal signal intensity to generate log2 relative copy-numbers, and removed probes corresponding to germline copy-number alterations(Gao *et al.*, 2022).

### Pre-processing of raw coverage data for downstream analysis (CCLE data)

The CCLE data was preprocessed in the same manner as the TCGA data. There were no samples with more than 5% autosomal loci with zero coverage, so no samples were excluded by this criterion. 703 probes out of 189915 were excluded because they had zero coverage in more than 40% of all samples. Two normal samples with questionable gender labels were excluded due to the balance of chrX and chrY signal intensities, bringing the total number of remaining samples to 324.

### TangentXY

Different approaches are applied to chrX and chrY in TangentXY. For chrX, we did the following:

1. The mean of the signal intensities for all loci of chrX in all male samples ( Μ_chrX male_) was calculated, as well as for the female samples (Μ_chrX female_). The chrX signal at each locus (i) of each male sample (j) was linearly transformed by the equation below. 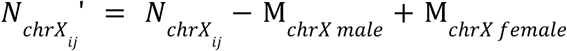
2. The portion of the noise space *N* corresponding to chrX of the male sample was replaced with *N_chrX_ij__*′ to produce *N’*.
3. We then performed SVD(Klema and Laub, 1980) on the portion of *N′* corresponding to the autosomes and chrX (*N_auto_*_+*c*ℎ*rX*_′), and retained the top *k* singular values (latent factors) from the resulting matrices to reconstruct *N_auto_*_+*c*ℎ*rX*_ʺ.
4. We performed Tangent normalization on the portion of tumor samples corresponding to autosomes and chrX (*T_auto_*_+*c*ℎ*rX*_) using *N_auto_*_+*c*ℎ*rX*_ʺ to yield their normalized log2 relative copy-numbers (*T_auto_*_+*chrX*_^′^).

For chrY,

1. We observed that signal differed between sequencing centers much more for chrY than other chromosomes, possibly because chrY has far fewer probes than other chromosomes (chrY: 402, median for other chromosomes: 6927). This batch variation significantly impaired subsequent analyses (data not shown). We therefore performed batch correction on the chrY signal. We set the average value of samples sequenced at the Broad Institute (BI), which had the largest sample size, as the baseline. We then subtracted the difference between each sequencing center’s average value and the BI average value from each sample. This ensured that the average value for each batch matched the BI average value. This approach is based on the same idea as the method described above for correcting for sex differences on chrX.
2. After the batch correction, the mean of the signal intensities for all loci of chrY were calculated for each male sample (μ_chrY_) and the median of the μ_chrY_ was calculated. The chrY signal at each locus (i) of each male sample (j) was linearly transformed by the equation below. 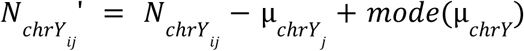
3. The portion of the noise space *N* corresponding to chrY of the male sample was replaced with *N_chrY_ij__*′ to produce *N’*.
4. We then performed SVD and Tangent normalization as we had for X and all other chromosomes, except that we restricted the normals to only males (*N_male_*′), to yield normalized log2 relative copy-numbers (*T_male_*′).
5. The normalized signal for chrY (*T_chrY_*′) was extracted and then reduced by 1 to be relative to the ploidy of each sample and then combined with *T_auto_*_+*chrX*_′ to obtain the final normalized value (T’).

### Performance evaluation of TangentXY, sex-matched Tangent, and Tangent

For the comparison of TangentXY with sex-matched Tangent or Tangent, 1000 pairs of normal and tumor samples, 500 each for females and males, were selected from TCGA. We attempted to include as many normal and tumor samples that were assayed in the same batch as possible, ending up with 1000 pairs of samples from 40 batches.

Performance was evaluated using signal and noise calculations as previously described(Gao *et al.*, 2022). In brief, we measured the ‘empirical signal’ as the standard deviation of median signal intensities among all chromosome arms and the ‘empirical noise’ as the median absolute difference between log2 relative copy-numbers of adjacent intervals (for sequencing data in which coverage was assessed by interval) or probes (for microarray data, in which signal intensities were assessed per probe).

### Segment level relative copy number estimation

To estimate the relative copy-number in each chromosome segment in a sample (tumor cells + non-tumor cells), the data normalized by TangentXY or Tangent were subjected to Circular Binary Segmentation (CBS)(Olshen *et al.*, 2004; Venkatraman and Olshen, 2007).

### Integer copy-number estimation of tumor cells

The relative copy-number of locus x (*R*(*x*)) in a sample, which is assumed to be a mixture of tumor cells and non-tumor cells, is defined as follows.

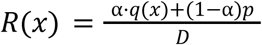

where *q*(*x*) is integer copy-number of tumor cell in the sample, *p* is the integer copy-number of non-tumor cell in the sample (i. e., 2 for autosomes and female chrX, and 1 for male chrX and chrY), α is the fraction of tumor cells in the sample (tumor purity). *D* is the ploidy of the sample, which is defined by:

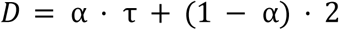

where τ is ploidy of the tumor cell. Consequently, the integer copy-number of the tumor cell in the locus x is obtained from the following equation.

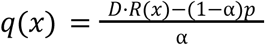

We used tumor purity and ploidy data called by ABSOLUTE(Carter *et al.*, 2012), which were downloaded from the GDC PanCanAtlas Publications page (TCGA_mastercalls. abs_tables_JSedit. fixed. txt) for α and τ. And we used the segment level relative copy number value generated by CBS for *R*(*x*) (see the above method).

### Classification of diploid and polyploid

To classify the TCGA samples as diploid and polyploid, we used purity and ploidy data called by ABSOLUTE(Carter *et al.*, 2012) (described above). For CCLE samples, we used purity and ploidy data (ccle_broad_2019_clinical_data. tsv) downloaded from cBioPortal. Samples with ploidy < 2. 5 and genome doubling = 0 were defined as diploid, and those with ploidy ≥ 2. 5 or genome doubling ≥ 1 were defined as polyploid.

### Classification of SCNAs in chrX and chrY

To classify SCNA in chrX and chrY, we took the following three steps: segment level classification, arm-level classification, and sample-level classification. First, we extracted the ploidy of each sample from the ABSOLUTE call. In the absence of copy number variations, the number of chrX in female cells is considered to be equal to the ploidy level, whereas in male samples, the number of both chrX and chrY is half the ploidy level. This number is denoted as *d* (ploidy in females, and ploidy/2 in males). The estimated integer copy-number less than *d* is considered copy number loss and greater than *d* is gain/amplification. We used thresholds of ±0. 2 based on the distribution of log2 integer copy numbers of chrX and chrY (data not shown) to assign SCNA classification to each segmentation. Since copy number = 0 cannot be expressed on a log2 scale, a threshold was set at the integer scale only for the Loss classification. The definitions for each classification are as follows.

– Loss: Integer CN < 0. 2 (Regardless of *d*)
– Subclonal loss: log2(0. 2) ≤ log2 integer CN < log2(*d*) - 0. 2
– Neutral: log2(*d*) - 0. 2 ≤ log2 integer CN < log2(*d*) + 0. 2
– Subclonal gain: log2(*d*) + 0. 2 ≤ log2 integer CN < log2(*d* + 1) - 0. 2
– Gain: log2(*d* + 1) - 0. 2 ≤ log2 integer CN < log2(*d* + 1) + 0. 2
– Amplified: log2(*d* + 1) + 0. 2 ≤ log2 integer CN

After assigning segment-level classification, arm-level classification was performed. If the total length of the segments classified as any of the above categories (Loss, Subclonal loss, Neutral, Subclonal gain, Gain, or Amplified) accounted for more than 50% of a chromosome arm, the arm was classified as the corresponding class. If no single class exceeded 50% of the total length, it was classified as Mixed.

Finally, a combination of p- and q-arm classifications resulted in chromosome-level labeling as follows:

– Loss: both of p arm and q arm are classified as loss
– Subclonal loss: both of p arm and q arm are classified as subclonal loss
– Neutral: both of p arm and q arm are classified as neutral
– Subclonal gain: both of p arm and q arm are classified as subclonal gain
– Gain: both of p arm and q arm are classified as gain
– Amplified: both of p arm and q arm are classified as amplified
– Mixed: all other combinations of p arm and q arm classifications

### ChrY loss call from Mitelman database

Data corresponding to each TCGA cancer type was extracted from the Mitelman database(Mitelman Database Chromosome Aberrations and Gene Fusions in Cancer), and the proportion of patients with karyotypes containing “-Y” was calculated. For comparison with the chrY deletion rate called by TangentXY, to avoid skewed deletion rates due to small sample sizes, comparisons were limited to cancer types with at least 10 samples in both the Mitelman database and TangentXY datasets.

### Association of SCNAs in sex chromosomes with gene mutations

Somatic mutation calls (mc3. v0. 2. 8. PUBLIC. LAML_PATCH. maf. oncokb. txt) were obtained from the GDC publications page (https://gdc.cancer.gov/about-data/publications/Pathways-2018). Only loss-of-function (LOF) mutations were analyzed. LOF mutations were defined as truncating events (frameshift deletions, frameshift insertions, nonsense, nonstop, splice site, and translation start site) or missense mutations predicted as deleterious by SIFT(Ng and Henikoff, 2002) or damaging by PolyPhen-2(Adzhubei *et al.*, 2013). The samples were divided according to the presence or absence of somatic point mutations in a gene and arm-level copy-number alteration in chrX/chrY (subclonal gain/gain/amplification or loss/subclonal loss). To evaluate the association between mutation status and copy number status while adjusting for potential confounders, we fitted a multivariable logistic regression model with copy number class as the dependent variable and mutation class as the primary independent variable. Tumor type, sex, and fraction of genome altered (FGA) were included as covariates. Finally, p-values were corrected for multiple testing using the Benjamini-Hochberg method(Benjamini and Hochberg, 1995). Only genes with the number of mutated samples greater than at least 1% of evaluable samples were included in the analysis.

### Survival analysis

Overall survival and progression-free survival data were downloaded from the GDC PanCanAtlas Publications page (TCGA-CDR-SupplementalTableS1.xlsx). Significance for Kaplan-Meier statistics was calculated with the log rank test using the R survival package(Therneau and Grambsch, 2013). Groups with either subclonal gain/gain/amplification or subclonal loss/loss were compared to groups without SCNAs at the whole chromosome level. To control for the difference in SCNA effects by tumor subtype, each tumor type was divided into subtypes based on a previous report(Ellrott *et al.*, 2025). Only tumor subtypes with at least ten samples in both groups were included. Results with a Bonferroni-corrected p-value < 0. 1 were considered statistically significant.

### Association of gene expression with SCNAs in chrX and autosomes

In this analysis, to investigate how gene expression changes with copy number change, the slope of individual gene expression relative to copy number was calculated using the following procedure. Only diploid tumor samples were used in this analysis. The gene expression file for each TCGA sample (“*. rna_seq. augmented_star_gene_counts. tsv”) was downloaded from the GDC Data Portal and the TPM values were extracted. For CCLE samples, we used a file downloaded from the DepMap portal (OmicsExpressionAllGenesTPMLogp1Profile. csv). The chrX genes were classified into inactive genes, escape genes, and variable genes based on data reported by Tukiainen et al(Tukiainen *et al.*, 2018), and only inactive and escape genes were included in the analysis. In addition, only genes with a median TPM greater than 1 were used in the analysis. In female samples, it was anticipated that inactive genes and escape genes would exhibit different behaviors, and that the slope of gene expression relative to copy number would differ in the regions where 1 < CN < 2 and 2 < CN, respectively (Fig. 4a). Therefore, the slope of gene expression relative to chromosome copy number was calculated for each gene type and region. For each individual gene, the relative gene expression levels (calculation method detailed below) and copy numbers across all samples were plotted, and the slope of the linear regression line was calculated. The expected gene expression level from a single chromosome was defined using the following criteria, and the relative gene expression level was calculated by dividing the gene expression value of each individual sample by this value. For the analysis of male samples, the median TPM value of the samples in the CN=1 region was defined as the gene expression per chromosome copy; for female samples, the median TPM value of the samples in the CN=2 region was defined as the per-copy expression because usually, only one of two copies of chrX is transcriptionally active for inactive genes. For escape genes, the gene is also expressed from Xi, but the expression level from Xi is known to be much lower than that from Xa(Carrel and Willard, 2005), so we used the same method to approximate the expression level per chromosome copy for escape genes as well as inactive genes. Furthermore, since expression levels per chromosome were expected to vary by carcinoma type, this operation was performed separately for each cancer type. The minimum number of samples required to draw a regression line was set to 10, and genes with fewer samples were excluded.

### Allele-specific gene expression analysis

GATK ASEReadCounter was used to generate allele-specific expression data. TCGA germline mutation data were downloaded from the GDC data portal (PCA. r1. TCGAbarcode. merge. tnSwapCorrected. 10389. vcf. gz). For CCLE samples, we called germline variants as described below. Biallelic SNPs were selected with GATK SelectVariants, and only heterozygous SNPs were further selected with BCFtools and used as input for ASEReadCounter. WES bam files were used as another input. The GATK LiftOverVCF pipeline was used to correct for differences in genome versions between the WES bam files and the mutation vcf files. Only ASE reads that mapped to regions corresponding to SNPs in the exons were extracted, based on the Ensembl Genome Browser annotations, and used in subsequent analyses. For each locus, the two allelic ratios that sum to 1 were calculated, and the larger of the two values was taken to be representative of the allelic ratio for that locus, which is referred to as “ASE value” in the text. The median ASE values for each chromosome in a sample were calculated to represent the allelicity for the chromosome in that sample.

### Estimation of cancer types exhibiting XCI dysregulation

We calculated the regression line between ASE values and tumor purity (Fig. S8d), and also calculated the difference between this line and each sample. Next, we used the Wilcoxon rank-sum test to compare the distributions of these values between a specific cancer type and all other cancer types, thereby identifying the tumor types prone to XCI dysregulation. After correction for multiple comparisons using the Bonferroni method, results with p-values < 0. 05 were considered statistically significant.

### Whole exome sequencing germline variant calls in CCLE samples

CCLE whole exome sequencing bam files were aligned to the hg38 reference genome. Variants were called with GATK (version 4. 2) HaplotypeCaller and were annotated with GATK CNNScoreVariants and filtered with GATK FilterVariantTranches with a truth sensitivity range of 99. 95 - 100. 00. The resulting variants were further filtered with Bcftools v1. 20 to remove all variants not annotated as PASS.

### Male bias in cancer incidence analysis

The same somatic mutation calls as described above were used. Duplicate mutations within tumors (from different matched normals) were removed, and when multiple tumors from the same patient carried the same mutation, only one was retained. Samples with unknown gender were excluded.

To identify EXITS genes—tumor suppressors that escape X inactivation—across all 11,000 TCGA tumors, only LOF mutations were analyzed. Tumor types enriched in one sex (PRAD, TGCT, BRCA, CESC, OV, UCEC, UCS) were excluded.

Sex bias was assessed using a permutation test. Male-to-female mutation ratios were calculated, and sex labels were randomly reassigned to mutations while preserving the total counts and sex-specific ratios within each cancer type. For each gene, p-values were derived from the number of permutations exceeding the observed male mutation count and corrected using the Benjamini–Hochberg method(Benjamini and Hochberg, 1995). A per–cancer-type analysis was also performed, in which one p-value was obtained for each gene–type pair, and FDR correction was applied across all such pairs. We detected a total of nine EXITS genes (Fig. S6).

For EXITS genes with chrY paralogs, genes located in PAR regions were excluded due to incomplete coverage and ambiguous read mapping. Associations between LOF mutations or copy loss in X-linked genes and copy loss of their chrY paralogs were assessed using Fisher’s exact test. Specifically, 2 x 2 contingency tables were constructed within each cancer type, comparing tumors with or without LOF events in the X-linked gene against those with or without chrY loss. For the pan-cancer analysis, p-values were combined across cancer types using Fisher’s method. For the per–cancer-type analysis, one p-value was obtained for each gene–type pair, and FDR correction was applied across all such pairs.

### Data Availability

TangentXY is available at https://github.com/beroukhim-lab/RtangentXY. All code for producing results and figures in this manuscript is available at https://github.com/beroukhim-lab/TangentXY_script.

## Results

### Complications in detecting copy-number changes in sex chromosomes

Whole-chromosome copy-number changes of chrX and chrY are not well-represented in The Cancer Genome Atlas (TCGA) copy-number data available at the NIH GDC Data Portal(Zack *et al.*, 2013; Taylor *et al.*, 2018; Sadagopan *et al.*, 2022). These data provide log2 copy-number ratios, which represent the log of the ratio between local copy-numbers and the genome-wide average. The mean log2 ratio for chrX in both male and female cancers is almost invariably 0, representing the genome-wide average copy-number, e. g. two in a diploid cell. Male tumors would be expected to usually have values of -1 (representing a 50% reduction in copy-number), and somatic variations would be expected to lead to non-zero values across many cancers.

These 0 values are a sign that signals reflecting whole-chromosome gains and losses of chrX are eliminated by overfitting to germline data from both male and female samples (Fig. 1a). An early step in the copy-number determination methods used by TCGA was to apply Tangent normalization to raw copy-number data obtained from either single nucleotide polymorphism (SNP) array signal intensities or variations in sequencing depth in whole-exome or -genome sequencing data. Tangent considers each copy-number profile to be a vector in an M-dimensional space, where M represents the number of loci at which copy-number estimates are derived (e. g. 1,852,600 dimensions for SNP 6. 0 data). Tangent then assumes that the subspace spanned by all copy-number data from normal control samples reflects variations that may be due to experimental noise, and thus removes this component of the copy-number profile from each tumor. In other words, it projects the data for each tumor *T_i_* onto the normal subspace *N*, and then subtracts this projection from *T_i_* to obtain a denoised copy-number profile (Fig. 1b). This approach is useful overall: In TCGA data, it improved signal-to-noise ratios across most of the genome by more than three-fold(Gao *et al.*, 2022). However, the normal subspace used by Tangent includes variations in chrX copy-numbers between males and females. Thus, this component of *T_i_* is removed, removing evidence of alterations in overall chrX copy-numbers. Moreover, although female tumors in which chrY contributes no signal retain the lack of chrY in the SCNA estimates, female tumors with some signal (e. g. due to noise) in some cases appear to have over two copies of chrY (Fig. 1a). Focal alterations within chrX and chrY are retained, as these are not present in normal samples.

**Figure 1.**
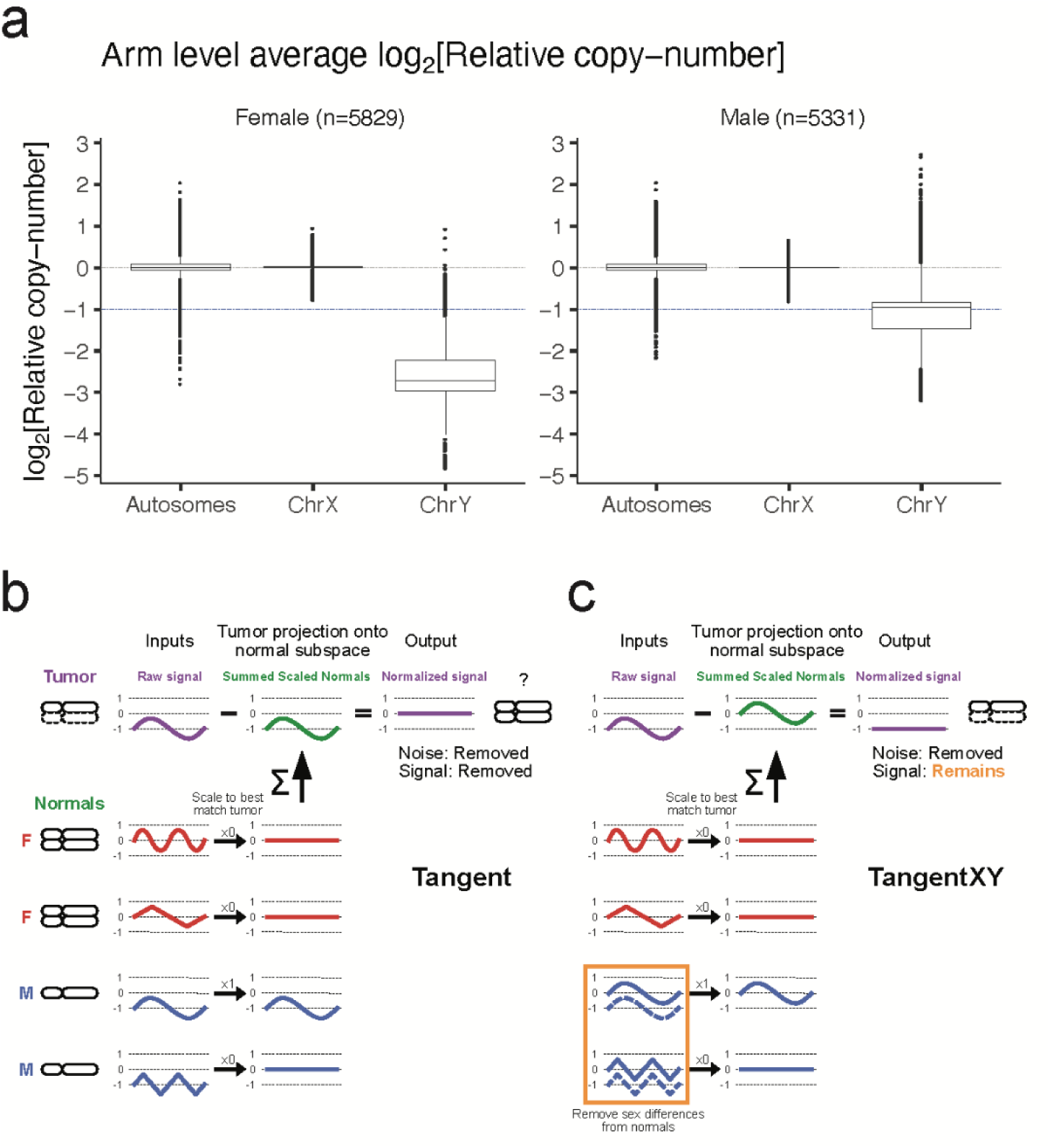
TangentXY addresses overfitting of chrX copy-number data in existing TCGA analyses, while retaining noise reduction capabilities. (**a**) Mean value of arm-level log_2_(Relative copy-number) per chromosome in segmentation files from SNP array data on 11160 tumor samples in the TCGA data portal (5829 females and 5331 males). The blue dashed line indicates the value chrX should have in males without chrX SCNAs. (**b, c)** Schematic overview of why chrX signal is lost in the current version of Tangent (**b**) and how it is overcome in TangentXY (**c**). Normalization of a female tumor sample with a single-copy deletion of chrX is shown as an example.

Therefore, we hypothesized that removing the sex differences from *N* would prevent overfitting (Fig. 1c). An alternative approach would be not to use Tangent normalization for sex chromosomes, but we were concerned that this approach would lose the noise-reduction benefits of Tangent. Similarly, normalizing against only sex-matched reference normals would reduce the available data on noise patterns in the dataset, especially in cases where few normals from a single sex are available (see below).

### TangentXY: a sex-informed approach to detecting copy-number alterations at sex chromosomes

We first evaluated those differences in whole exome sequencing (WES) data from 10,441 normal samples and 9,854 tumor samples across 33 tumor types from TCGA that met quality-control metrics. As expected, signal distributions for chrX in normal female and male samples are near 0 (and similar to the autosomes) and -1, respectively (Fig. S1a). Therefore we expected that shifting the male chrX signals to the same levels as the female chrX would remove sex differences from the normal subspace. For chrY, we found that the signal distribution is very different from that of chrX (Fig. S1c). Since females do not have chrY, their signals contain a large number of zeros in the raw coverage data and do not reflect the noise distribution in males (Fig. S1c).

ChrY exhibited more variation in peak signal intensities than we had expected, even in male samples. We hypothesized that this reflects mosaic loss of chrY in many of the reference “normal” male samples. Mosaic loss of chrY is observed frequently as men age, including in >40% of blood samples from healthy men over 70 years old(Thompson *et al.*, 2019). Normalizing a cancer sample against a reference sample with mosaic loss of chrY would lead to an artifactual appearance of a gain in chrY copy number.

Therefore, we designed TangentXY, an extension of Tangent that accounts for sex differences in chrX and mosaic loss of chrY. TangentXY performs a linear transformation on chrX signals in male samples within the normal reference set so that they have the same mean as females within that reference set (see Methods). TangentXY only uses chrY data from males in the reference set for normalization. We also designed TangentXY to perform its linear transformations using only a subset of the dimensions offered by the normal samples. As one adds normal samples to the TangentXY inputs, the number of dimensions in the normal subspace increases–eventually raising concerns that true signals in the copy-number profiles will be removed due to overfitting. Therefore, TangentXY performs singular value decomposition (SVD) on the entire normal subspace (excluding chrY loci) to rank dimensions (aka “latent factors”) by their contribution to signal intensity profiles (represented as their singular values). TangentXY then reconstructs the normal subspace with a subset of the highest-ranking latent factors, with the number of latent factors set by the user. We ran TangentXY with varying numbers of latent factors, and found that signal-to-noise ratios (estimated as in prior studies(Beroukhim *et al.*, 2007, 2010; Gao *et al.*, 2022)) were maximized at ∼100 latent factors; at higher numbers, the amount of signal began to erode (Figs 2a-c).

**Figure 2.**
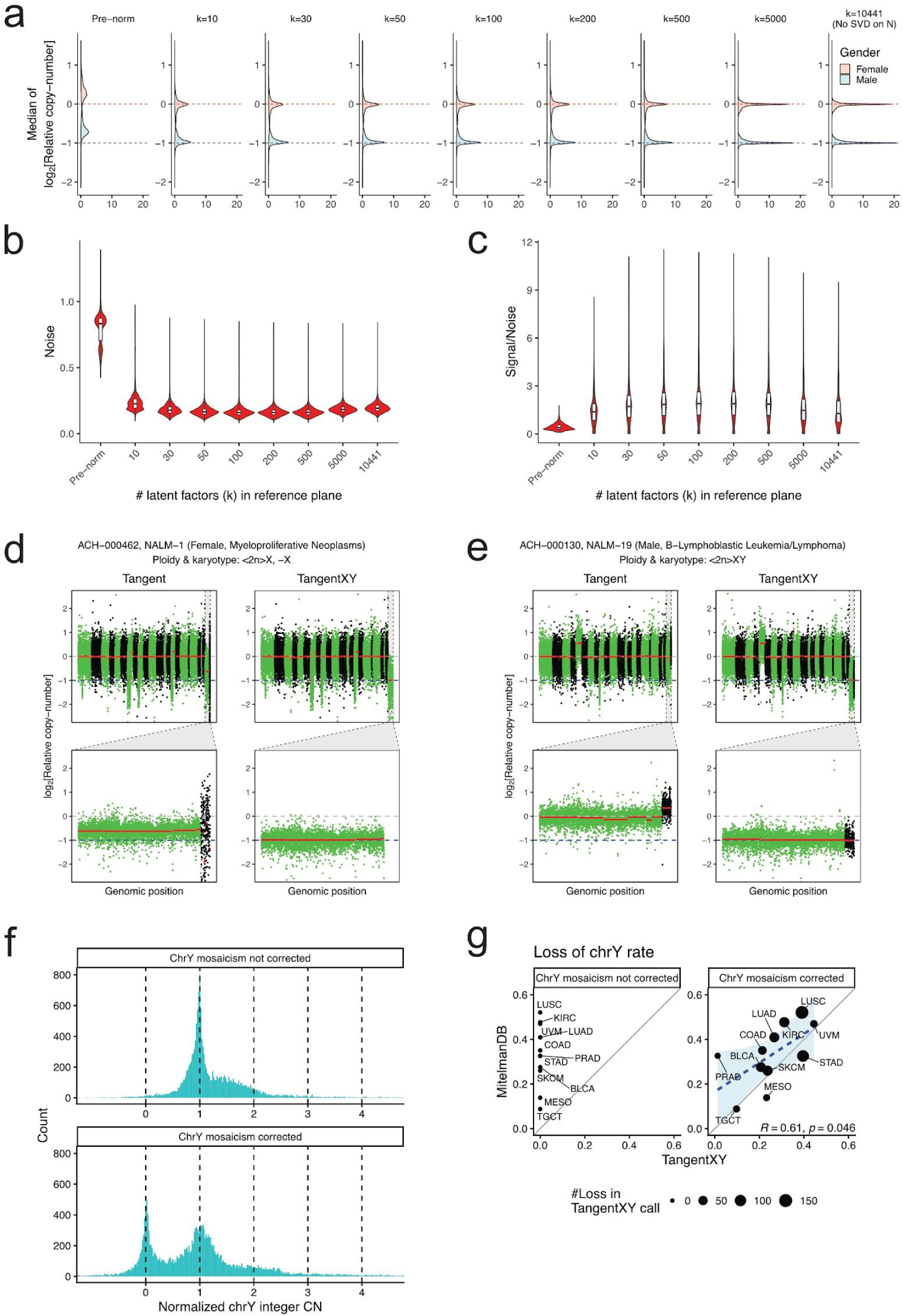
Performance of TangentXY. (**a**) Distribution of median log_2_(Relative copy-number) of chrX before and after normalization with TangentXY using reconstructed normal subspaces with different numbers of latent factors. (**b, c**) Noise levels (**b**) and signal-to-noise ratios (**c**) for pre- and post-TangentXY data. (**d, e**) Representative images of Tangent/TangentXY normalization on CCLE cell lines with validated karyotype annotations. Top panels: Entire genome (green, odd-numbered chromosomes and chrX; black, even-numbered chromosomes and chrY); bottom panels: enlarged view of chrX (green) and chrY (black). Red lines indicate estimated copy-number for each segment after CBS. (**f**) Histogram of chrY copy number in male tumors before correcting for chrY mosaicism (upper panel) and after correcting for chrY mosaicism (lower panel). (**g**) Fraction of clonal loss of chrY in Tangent XY call and the Mitelman database

TangentXY performs better than simply normalizing against sex-matched controls. We compared noise reduction performance between sex-matched Tangent and TangentXY using various numbers of normal control samples across both sexes. Whether using control cohorts of various sizes that were evenly divided by sex (Fig. S2a), or cohorts of 500 normal controls that were unevenly divided between sexes (Fig. S2b), the noise reduction of TangentXY was greater than sex-matched Tangent in all cases, with the greatest benefits when few sex-matched controls were available. We also found that the noise reduction performance for autosomes was almost identical between the former version of Tangent and TangentXY (Fig. S3a), though possibly favoring TangentXY when the control datasets were large in terms of signal-to-noise ratio (Fig. S3b).

We validated TangentXY output on copy-number data from the Cancer Cell Line Encyclopedia (CCLE) (Ghandi *et al.*, 2019) in 130 cancer cell lines with experimentally validated karyotype annotations. We ran circular binary segmentation (CBS) after Tangent or TangentXY normalization, and then manually reviewed both. Whereas Tangent normalization generally detected little whole-chromosome copy-number deviation from the euploid level for chrX and chrY, TangentXY recapitulated copy-number changes in chrX and chrY indicated in the karyotyping data (Fig. 2d-e, Fig. S4a-i). TangentXY showed significant improvements in accuracy, sensitivity, and specificity across all alteration types (although copy number gains were extremely rare on chrX and were not observed on chrY in this dataset) (Fig. S4j). This improvement was primarily observed in male samples, which is expected because of the single copy of sex chromosomes in males.

### ChrY mosaicism in normals contaminates somatic copy-number estimates

However, we found that this approach was insufficient for normalizing chrY. ChrY signal intensities do not only differ between normal male and female tissues; even in males, normal samples exhibited signal intensities in chrY that were much more variable relative to signal intensities in chrX or any other chromosome (Fig. S1c). When we used these raw signal intensities as reference data for TangentXY, we detected a spread of somatic copy-number estimates for chrY that also differed from copy-number profiles across the rest of the genome, with far more apparently subclonal gains, with very few clonal events (Fig. 2f, upper panel).

We suspect that these findings reflect frequent copy-number mosaicism of chrY in normal tissue(Jacobs *et al.*, 1961, 1963; Pierre and Hoagland, 1972). This mosaicism will contaminate somatic copy-number estimates from chrY in a similar way to how differences in chrX dosage between normal male and female samples contaminated somatic copy-number estimates from chrX when raw chrX data were used for normalization (Fig. 1b). For this reason, we desired to adjust the chrY signal intensities from normal samples to reflect the best estimate of “pure” tissues in which chrY mosaicism does not exist. We assumed that the largest plurality of normal male tissues had little mosaicism, so that the modal level chrY across all male normal tissues would reflect this “pure” non-mosaic state. Under this assumption, we adjusted the signal intensity for each normal male reference sample to equal the mode of these averages across all males (see Methods), and used these adjusted signal intensities in TangentXY.

This adjustment appeared to vastly improve the somatic chrY copy-number estimates produced by TangentXY (Fig. 2f, lower panel). As with other genomic loci, we now observe clear “peaks” in somatic copy-numbers at 0 and 1 copies, with a hint of a peak at 2 copies. We calculated the fraction of cancers with clonal loss of chrY. These fractions were much closer matches to prior estimates of clonal chrY loss based upon fluorescence in-situ hybridization (FISH) and spectral karyotyping studies(Mitelman Database Chromosome Aberrations and Gene Fusions in Cancer; Wang and LaFramboise, 2019) compared to copy-number estimates prior to our adjustments of chrY signal intensities in the normal reference data (Fig. 2g). However, significant discrepancies were observed for prostate adenocarcinoma (PRAD). For PRAD, a validation study with a much larger sample size (n=2053) than the Mitelman database reported very low frequency of loss of chrY (0. 5%), which is consistent with TangentXY’s estimates(Stahl *et al.*, 2012).

### SCNAs in chrX and chrY across tumor types

We next applied TangentXY to all 9,854 tumor samples with WES data from TCGA, spanning 33 tumor types (Fig 3). Testicular germ cell tumors (TGCT) and colorectal cancer types (COAD and READ) showed the highest rate of chrX gain/amplification, which is consistent with previous reports (Fig. 3a)<u>(Kraggerud *et al.* 2002; Dutrillaux *et al.* 1986; Muleris *et al.* 1990)</u>. The frequency of whole chrX gain/amplification varied by tumor type, but males were generally more likely to obtain gain/amplification than females within the same tumor type (Fig. 3a). On the other hand, partial chrX deletion (either hemizygous or subclonal) was observed much more frequently in female samples than in males and complete (homozygous) loss was not observed in either sex, reflecting the fact that chrX has genes that are essential for cell survival. Although a few male samples were also found to have subclonal loss, most of them were observed only in cells that experienced genome doubling, suggesting that such cells do not completely lose chrX even after the deletion (Fig. 3b). The reason why partial chrX deletions are more common in females than males is likely similar: males have only one copy of chrX, while females have two.

**Figure 3.**
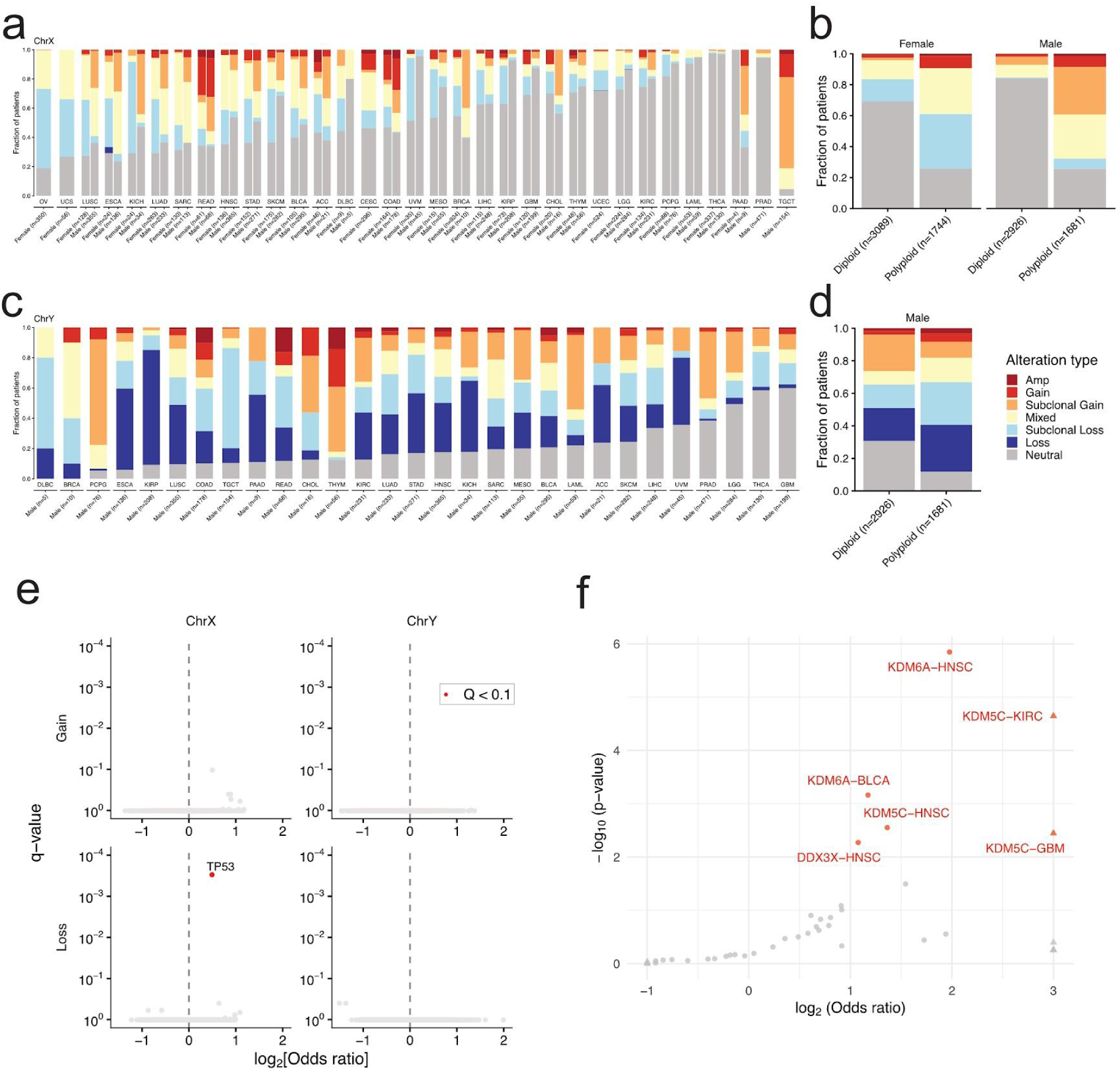
Associations between sex chromosome SCNAs and other features of the tumors in which they occur. (**a-d**) Fraction of tumors with whole-chromosome and whole-arm SCNAs in chrX and chrY across tumor type (**a, b**) and ploidy class (**c, d**). When both p and q arms are Amp, Gain, Subclonal Gain, Neutral, Subclonal Loss, or Loss, the samples are labeled respectively; otherwise labeled as Mixed. (**e**) Association of SCNAs in sex chromosomes with gene mutation. Genes with Q-value < 0. 1 are shown in red. P-values were calculated using logistic regression adjusted for tumor type, sex, and FGA. (**f**) Association of LOY and mutations in EXITS genes across tumor types. Significant associations are shown in red. Odds ratios falling outside of [-1,3] range were capped to improve readability and the corresponding gene-type pairs were marked with triangles.

Whole chrY deletion was most frequent in kidney renal papillary cell carcinoma (KIRP) (Fig. 3c). Compared to chrX, chrY showed higher rates of both amplification and clonal loss, suggesting that these alterations of chrY are better tolerated.

We next assessed the association between gene mutation and SCNAs in sex chromosomes. TP53 was the only gene that showed a highly significant association with losses of chrX (as previously noted (Taylor *et al.*, 2018)) (Fig. 3e, Table. S1). However, there was no significant association with chrY alterations.

### SCNAs in chrX and chrY predict prognosis of some cancers

We performed a survival analysis to see whether amplification or deletion of sex chromosomes is associated with overall survival (OS) or progression-free survival (PFS). We performed the analysis separately for males and females and separated tumors by cancer subtype (eg endometrioid vs serous endometrial cancers) because of differing biology between cancer subtypes(Ellrott *et al.*, 2025) and differing genetics that can confound association analyses(Carrot-Zhang *et al.*, 2020). As a result, we identified one significant association with OS (Fig. S5a) and two with PFS (Fig. S5b-c). In BLCA_1, a basal squamous subtype of BLCA, patients with chrY amplification had longer overall survival (OS) and progression-free survival (PFS) than those without. Interestingly, while it has been reported that chrY loss is associated with a poor prognosis in BLCA<u>(Abdel-Hafiz *et al.* 2023; Chen *et al.* 2025)</u>, association between chrY amplification and a favorable prognosis has not been reported. In the male patient cohort of LUAD_3, chrX amplification also appeared to be associated with shorter PFS.

### ChrY loss may account for frequent mutations of KDM5C and KDM6A in male cancers

We previously noted that *KDM5C* and *KDM6A* are among the genes mutated more frequently in male cancers compared to female cancers(Dunford *et al.*, 2017). Both genes are tumor suppressor genes located on chrX that escape X inactivation, and we hypothesized that their higher mutation rates in male cancers might contribute to the male bias in cancer incidence. However, these two genes have homologs on chrY (*KDM5D* and *UTY*, respectively), raising the question why these homologs do not suppress cancers in males. We hypothesized that male tumors undergo chrY loss at rates that exceed chrX loss in females (Figs. 3c-d), resulting in higher rates of biallelic inactivation of *KDM5C/D* and *KDM6A/UTY*, respectively. However, this hypothesis would predict an association between mutations in KDM5C or KDM6A and chrY loss–an association we did not observe.

Using TangentXY, we now observe a significant association between mutations in *KDM5C* and *KDM6A* and chrY loss across all male tumors in TCGA (p=10^-5^ and 3x10^-4^, respectively), and specifically in head and neck squamous cell carcinomas (HNSC), bladder carcinomas (BLCA) for KDM6A (FDR q=6x10^-5^ and 0. 009, respectively), and kidney renal clear cell carcinoma (KIRC), HNSC, and glioblastoma multiforme (GBM) for KDM5C (FDR q=5x10^-4^, 0. 03, and 0. 03, respectively, Fig. 3f). We were also able to detect associations between chrY loss and mutations in *DDX3X* in HNSC (FDR q=0. 04). We were now able to detect these significant associations because we had improved the accuracy of chrY loss estimates by accounting for chrY mosaicism in the normal male tissue reference samples. We conclude that higher rates of cancers driven by *KDM5C* and *KDM6A* in males are due to higher rates of chrY loss in males relative to chrX loss in females.

### Amplified chrX is evenly distributed between Xa and Xi in females

We next asked whether amplified copies of chrX were transcriptionally active. In normal female cells, X chromosome inactivation (XCI) results in one of the two chrX becoming inactive X chromosome (Xi) and the other becoming active X chromosome (Xa)(Lyon, 1961). In Xi, most genes are “inactive” (not expressed), whereas “escape genes” retain transcriptional activity. A third category of “variable genes” varies across cell types(Berletch *et al.*, 2011). Even escape genes, however, have lower expression in Xi than Xa(Carrel and Willard, 2005). All genes in Xa remain transcriptionally active. The single copy of chrX in males is always Xa. A prior study that compared shapes of Xi and Xa found that the amplified chrX in both male and female cancers always resembles Xa(Dutrillaux *et al.*, 1986; Muleris *et al.*, 1990), whereas a more recent study evaluating expression levels found XCI occurs in some amplified copies of chrX in male cancers(Sadagopan *et al.*, 2022).

We examined expression levels of both inactive and escape genes in both female and male cancers. If amplified chrX is transcriptionally active, we would expect a single additional copy of chrX to double the expression of chrX genes, and further additional copies to add equally to gene expression (slope=1, Fig. 4a). In contrast, if amplified X is transcriptionally inactive, we would anticipate no increase in expression of inactive genes, and small increases in expression of escape genes (slope<1, Fig. 4a). With an even mix of Xa and Xi, we would expect the slope to average 0. 5 for inactive genes and escape genes in males, and slightly higher than 0. 5 for escape genes in females.

**Figure 4.**
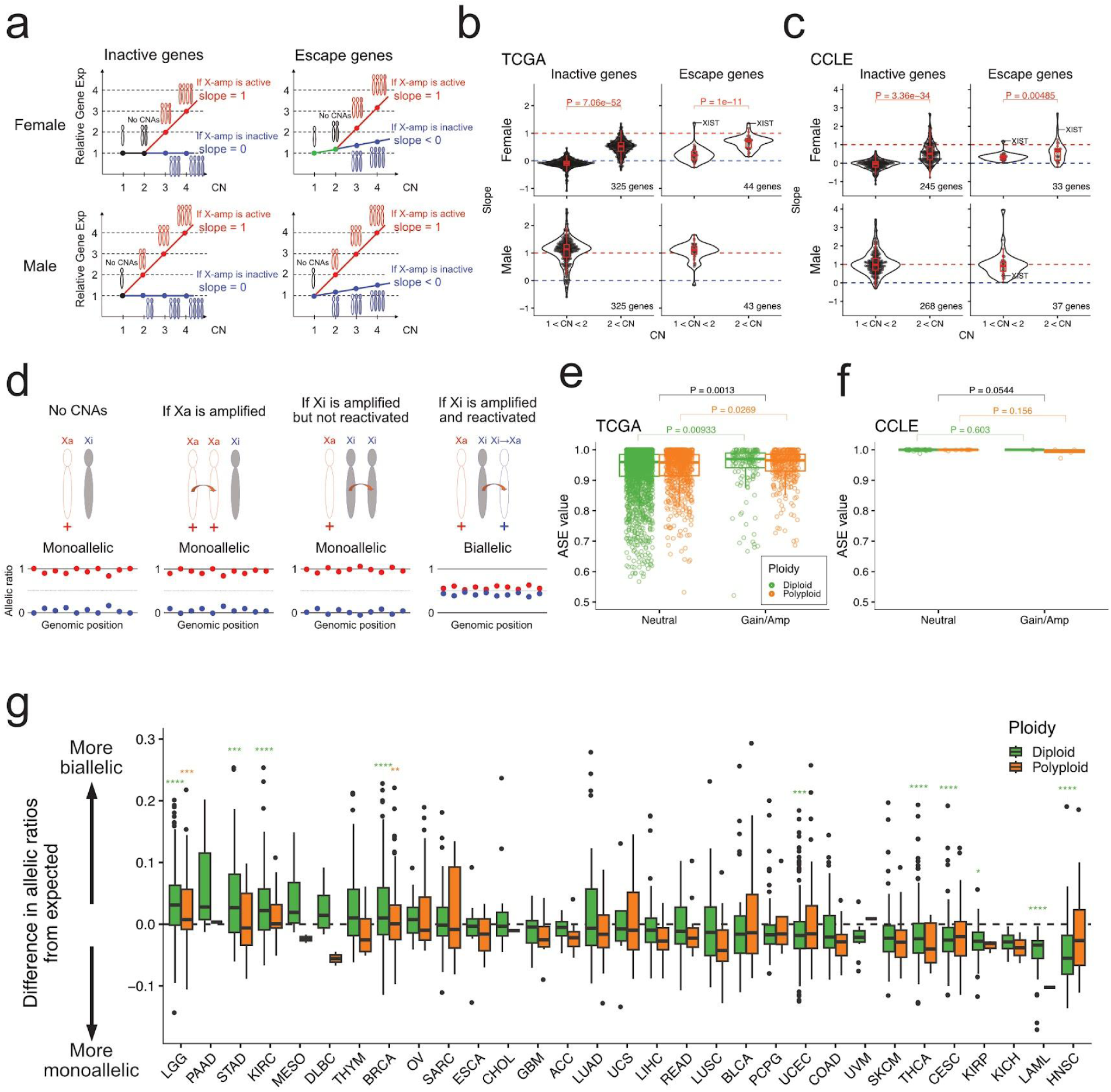
Characteristics of chrX regulation in aneuploid tumor cells. (**a**) Schematics of expected chrX gene expression patterns in different scenarios. The slope of gene expression with respect to chrX amplification changes depending on whether Xa or Xi is amplified. In female samples, the slope is expected to be different between copy-number less than 2 vs. greater than 2 because one of the two copies of chrX is inactive. (**b, c**) Slopes of inactive and escape genes for each copy-number region in TCGA samples (**b**) and CCLE cell lines (**c**). (**d**) Schematics of ASE ratio in different scenarios. The higher value (> 0. 5) was used as the ASE value in the downstream analyses. (**e, f**) Median ASE values for samples with no arm-level alteration and ones with whole amplification in chrX in TCGA samples (**e**) and CCLE cell lines (**f**). Plots for TCGA samples segregated by tumor type are shown in **Supplementary Figure 8a**. (**g**) The difference between the median ASE value for each sample and the regression line between median ASE values and tumor purity for each cancer type and ploidy class. The larger this value, the greater the biallelicity not explained by contamination with normal cells, i. e., the stronger the suspicion that XCI is occurring. P-values were determined by a Wilcoxon rank-sum test and adjusted by Bonferroni method (*, p < 0. 05; **, p < 0. 01; ***, p < 0. 001; ****, p < 0. 0001).

The data indicated that amplifications of chrX are not biased towards either Xa or Xi in females, and that amplified chrX always remains active in males. The average slope for inactive genes in females with chrX gains (CN>2) was 0. 51–statistically no different than 0. 5, whereas the average slope for escape genes was significantly higher at 0. 67 (p=2. 02x10^-5^). In males with chrX amplifications (CN>1), the average slope of both inactive and escape genes was approximately equal (p=0. 58) at 1. 05 and 1. 05 respectively (Fig. 4b). Analysis of copy-number data from 288 cell lines in the Cancer Cell Encyclopedia (CCLE) gave similar results (Fig. 4c). A similar analysis for autosomes yields a slope of approximately 1 (Fig. S7).

### Amplified Xi is not reactivated

We also examined whether Xi reactivation can occur in female cancer cells by performing allele-specific expression (ASE) analysis, in which we consider the higher of the two possible allelic ratios for transcripts of each gene (Fig. 4d). If Xi is never reactivated, in pure tumors we would expect uniform monoallelic expression for inactive genes on chrX in all cases, with ASE values near one. If Xi is reactivated, ASE values would decline towards 0. 5 (balanced expression between the two alleles; Fig. 4d). In tumors with substantial proportions of contaminating normal cells, ASE values would be reduced towards 0. 5 even when the tumor cells exhibited monoallelic expression. However, we would expect ASE values to remain higher when Xa is gained (including due to whole-genome doubling resulting in polyploidy) because a larger fraction of chrX in the sample would reflect Xa.

Indeed, we found that the ASE values approached one in almost all of the chrX amplified samples and were significantly higher in samples with chrX amplification compared to those without (p=0. 0013, Fig. 4e). This was true for both diploid and polyploid samples. The median ASE value was around 0. 5 for autosomes, which usually exhibit biallelic expression, confirming the accuracy of ASE analysis (Fig. S8b). Therefore we exclude the possibility of prevalent Xi reactivation after amplification. We further investigated whether this is because Xi is not being amplified or whether it is being amplified but simply not reactivated. To this end, we examined not only ASE but also the allele frequency (AF) of the DNA. Typically, in diploid cells with no copy number alterations, the DNA AF corresponding to ASE is expected to be around 0. 5 (with one copy each of Xa and Xi), and this was indeed the case (although a few samples showed a slight bias toward 1, most of these were due to loss of heterozygosity (LOH) and were considered reasonable) (Fig. S8c, left). In contrast, in samples with chrX gain (with one additional copy of the chrX), the DNA AF should theoretically be 0. 67 (i. e. two Xa out of three chrX); conversely, if Xi is amplified, it is expected to be 0. 33 (= one Xa out of three chrX). Clusters were observed near both theoretical values, confirming that there are cases where Xi is amplified in the same way as Xa (Fig. S8c, center). Similarly, in the samples with chrX amplification (where two or more copies were added), we confirmed that peaks were present near DNA AF of 0. 75 (three Xa and one Xi), 0. 5 (two Xa and two Xi), and 0. 25 (one Xa and three Xi) (Fig. S8c, right). Taken together, although Xi can be amplified as well as Xa, even when amplified, it is rarely reactivated.

Overall, low ASE values tended to be observed in impure tumors (Fig. S8d), as expected. Indeed, among CCLE cell lines, which should have no contaminating normal cells, we observed almost no deviation from monoallelic expression in chrX (Fig. 4f).

XCI dysregulation has previously been described in ovarian, lung, and breast cancers(Kristiansen *et al.*, 2005; Li *et al.*, 2006; Chaligné and Heard, 2014; Winham *et al.*, 2019). As described above, we observed the expected relationship between ASE value and tumor purity (Fig. S8d); however, samples deviating from the regression line were also observed sporadically, suggesting occasional XCI dysregulation. Therefore, we calculated the difference between each sample’s ASE value and the regression line and considered samples with a positive difference to be those exhibiting XCI dysregulation. In TCGA samples, we did observe a statistically significant deviation towards biallelic expression in low-grade gliomas, stomach adenocarcinomas, kidney cancers, and breast cancers after regressing out effects of tumor impurity (Fig. 4g). However, ASE values went below 0. 6 in only ten cases in the entire dataset (Fig. 4e and Fig. S8d), whereas ASE values in autosomes–which do exhibit biallelic expression–were almost always below 0. 6 (Fig. S8b). We conclude that the amplified copy of chrX rarely undergoes reactivation in cancer, if ever.

## Discussion

The detection of SCNAs in sex chromosomes of cancer cells has been hampered by intrinsic sex differences in copy-number. We show that TangentXY recapitulates copy-number changes of chrX and chrY while reducing noise. We also found that gains of chrX in females are equally likely to be the active or inactive copies and found little evidence for the reactivation of chrX in any cancer type.

Several CNV inference methods employing similar concepts to Tangent normalization have been reported. Examples include comunee 2. 0, Rawcopy and dry-clean<u>(Daenekas *et al.* 2024; Mayrhofer *et al.* 2016; Deshpande *et al.* 2019)</u>. Similar approaches to those we take here to extend Tangent to sex chromosomes might also be applied to these other methods.

TangentXY is mostly parameter-free, except for the number of latent factors (*k*) for reconstructing *N* in SVD. To determine the optimal *k*, one can re-run TangentXY using varying *k* values to maximize noise reduction while avoiding signal loss in the normalized chrX. In practice, however, the differences in noise reduction between different values of *k* tend to be small when *k* is greater than 30, and loss of signal due to overfitting tends to be small with *k* values less than 500–leaving a wide range of nearly-equivalent *k* values to choose from. For chrY, normalization was achieved by using only male normal samples as references. Given that the performance of Tangent normalization improves with the number of reference normal samples, we would like to take advantage of the female samples. However, the signal distribution of the female samples, which do not have chrY, is so different from that of male samples that it renders them unuseful and even counterproductive.

Using the newly developed TangentXY, we determined the representation of each SCNA type in sex chromosomes across 33 cancer types. Interestingly, we found that chrX amplification was more common in male samples than in female samples for almost all tumor types (Fig. 3a). Although there are few publications on this phenomenon, it has been reported that chrX amplification is highly prevalent in male breast cancer patients and that the androgen receptor located on the q-arm is important for neoplastic transformation(Di Oto *et al.*, 2015, 2018). In our study, the number of male breast cancer patients was small (10 cases), but half of them had whole/arm-level chrX amplification (Fig. 3a).

For chrY, we found more copy-number deletions than copy-number amplifications across cancer types (Fig. 3b), suggesting that chrY is not essential for cell survival, but rather that the deletion of chrY may favor cancer cell growth. It was recently reported that deletion of the chrY contributes to cancer cell growth by suppressing cancer immunity in patients with bladder cancer(Abdel-Hafiz *et al.*, 2023).

Based on the relationship between gene copy-number and expression of chrX, we found that the amplification of chrX occurred equally in Xa and Xi, and that, when Xi was amplified, it underwent reactivation only rarely, if ever, across TCGA samples. In this analysis, some samples without copy-number change in chrX showed skewed allelicity, however. This might reflect disruption of non-amplified XCI in some tumor types(Kristiansen *et al.*, 2005; Li *et al.*, 2006; Chaligné and Heard, 2014) or the effects of contaminating normal cells that had inactivated the other X allele.

In detecting SCNAs, varying allelic ratios across the genome can serve as an alternative signal to sequencing depth(Myers *et al.*, 2024). In principle, this could be effective in chrX in female samples. However, male samples will always be monoallelic in chrX and thus require analyses of sequencing depth. Across the genome, we anticipate that the optimal results will be obtained by combining both allelic ratios and sequencing depth, using methods to normalize sequencing depth such as TangentXY, as described here.

## Acknowledgments

This work was supported by the Gray Matters Brain Cancer Foundation, Pediatric Brain Tumor Foundation, Fund for Innovative Cancer Informatics, Team Jack Foundation and Break Through Cancer. We thank Srinivas R. Viswanathan, Cheng-Zhong Zhang, Alison M. Taylor, and Esther Rheinbay for their valuable discussions.

---

**Supplementary Table 1.** Associations between sex chromosome SCNAs and gene mutations.

**Supplementary Table 2.** Associations between sex chromosome SCNAs and progression-free survival (PFS) in each tumor type.

**Supplementary Figure 1.**
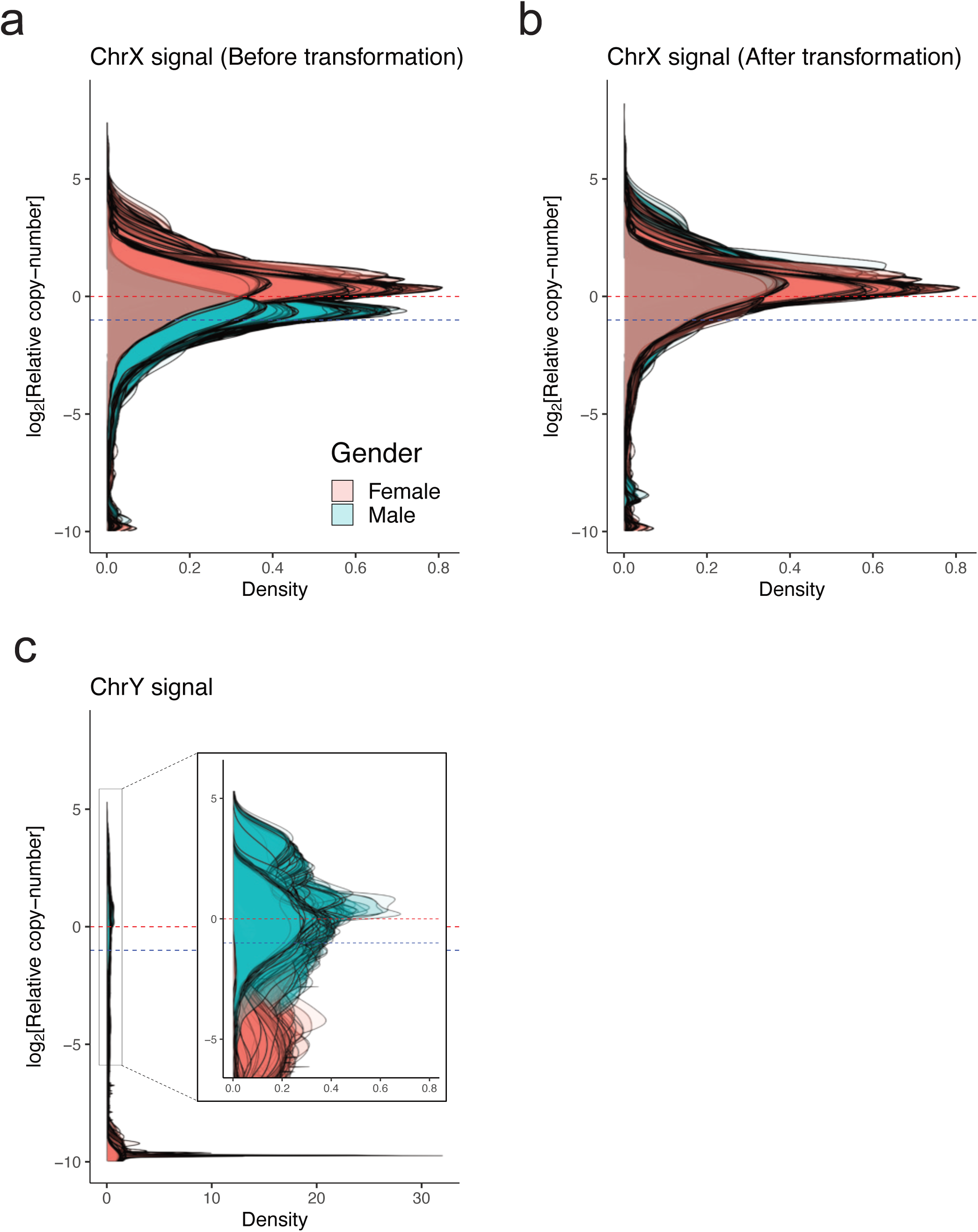
Signal distribution of WES coverage data in TCGA normal samples. (**a**) ChrX signal distribution before linear transformation. (**b**) ChrX signal distribution after linear transformation. (**c**) ChrY signal distribution.

**Supplementary Figure 2.**
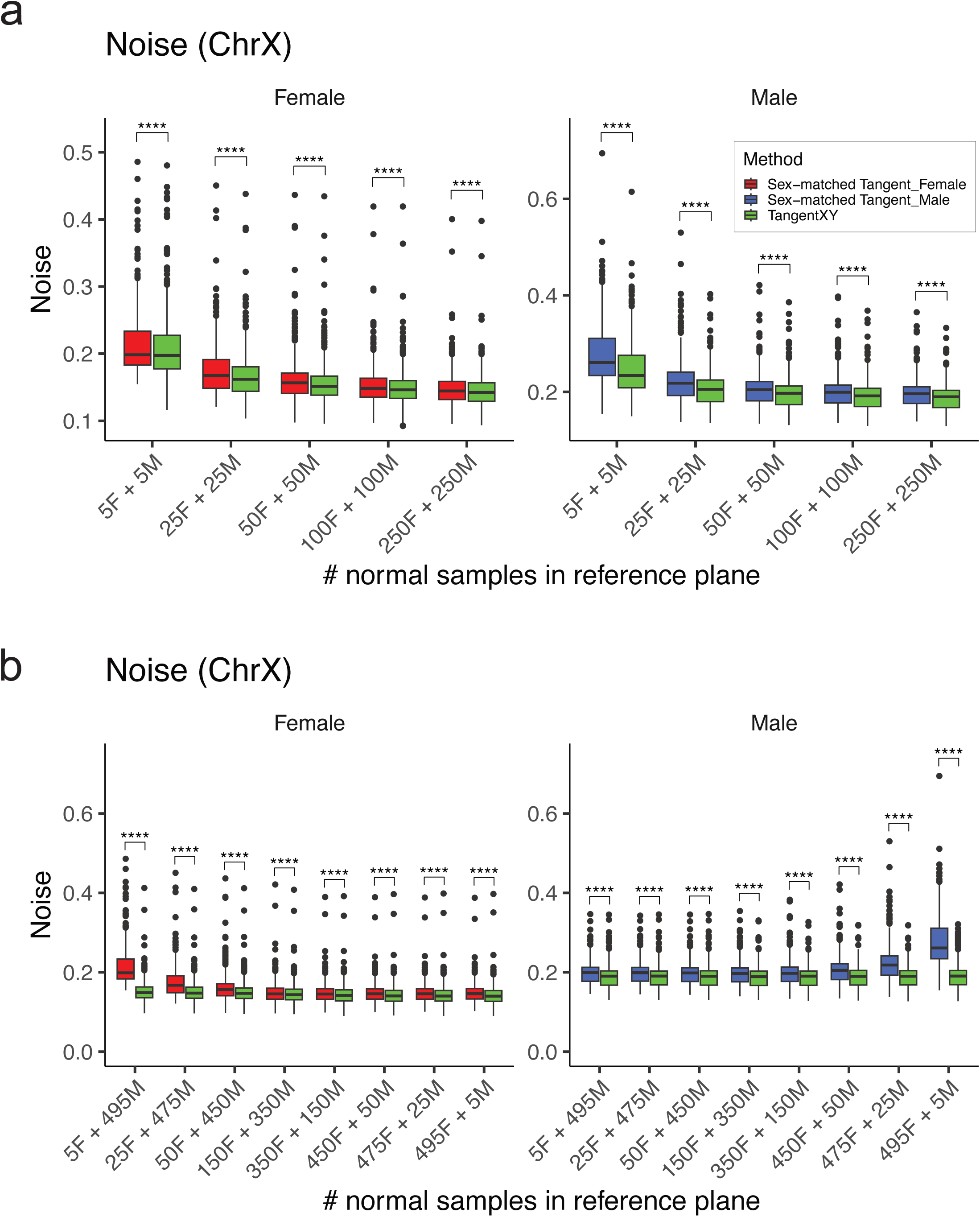
Noise reduction performance of TangentXY and sex-matched Tangent on chrX. (**a**) Box plots indicate chrX noise levels for pre-normalization, TangentXY, and sex-matched Tangent with different numbers of reference normal samples, equally divided between males and females. (**b**) Box plots indicate chrX noise levels for TangentXY and sex-matched Tangent using 500 normal samples with various female/male ratios. P-values were determined by Wilcoxon signed-rank test and adjusted by Bonferroni method (****, p < 0. 0001).

**Supplementary Figure 3.**
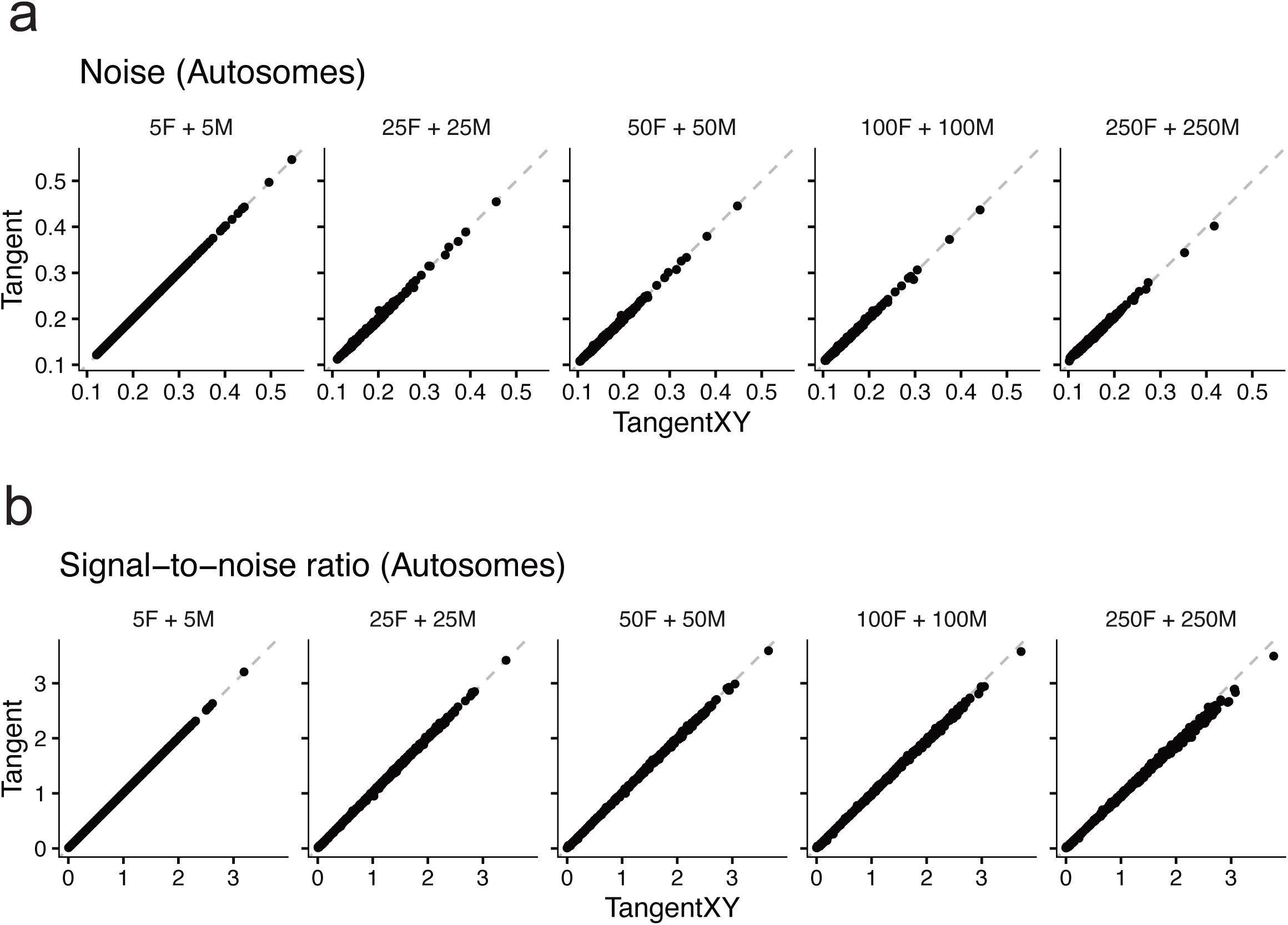
Noise reduction performance of TangentXY and Tangent on autosomes. Scatter plots indicate Tangent versus TangentXY noise levels (**a**) and signal-to-noise ratios (**b**) on autosomes.

**Supplementary Figure 4.**
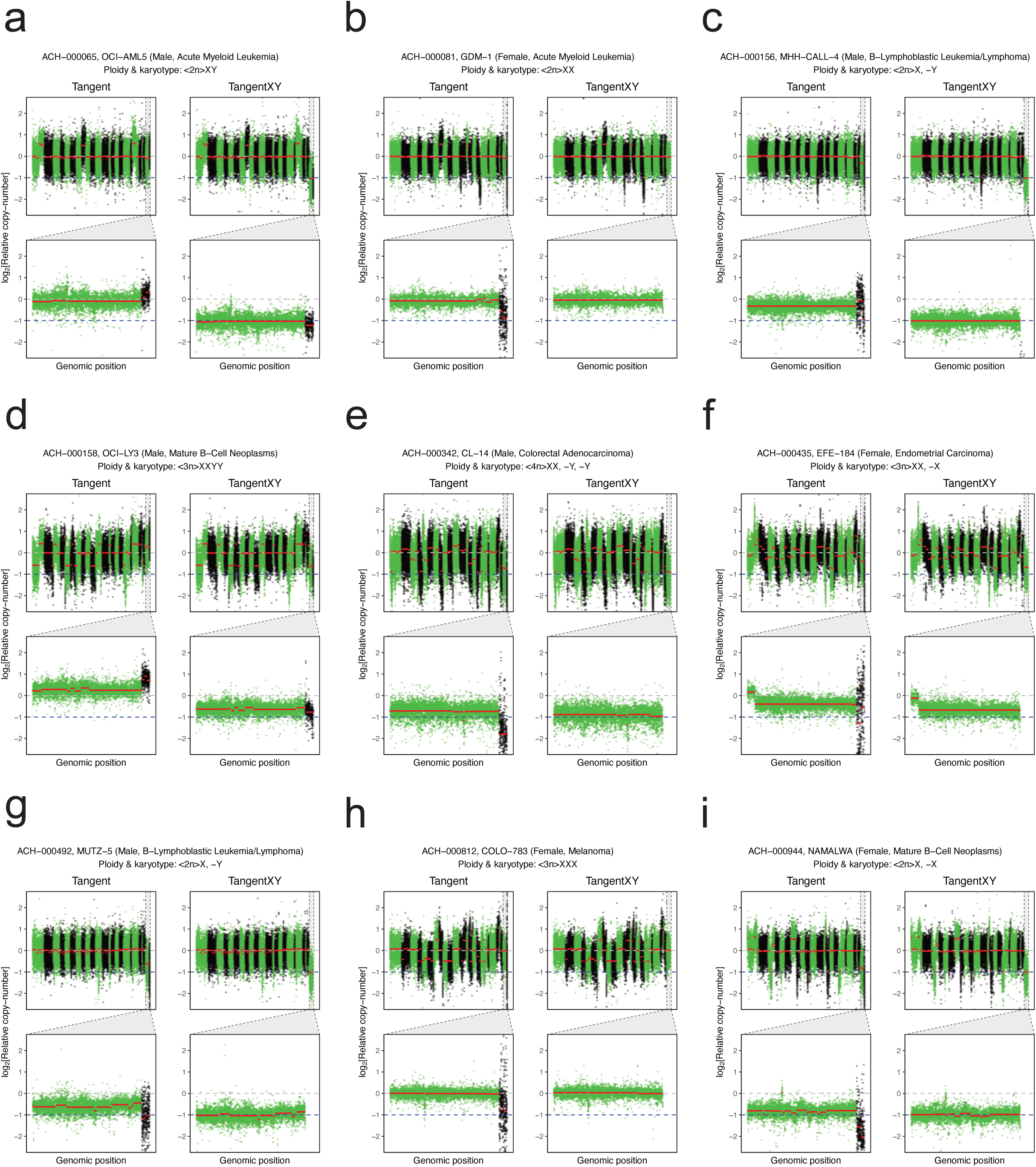
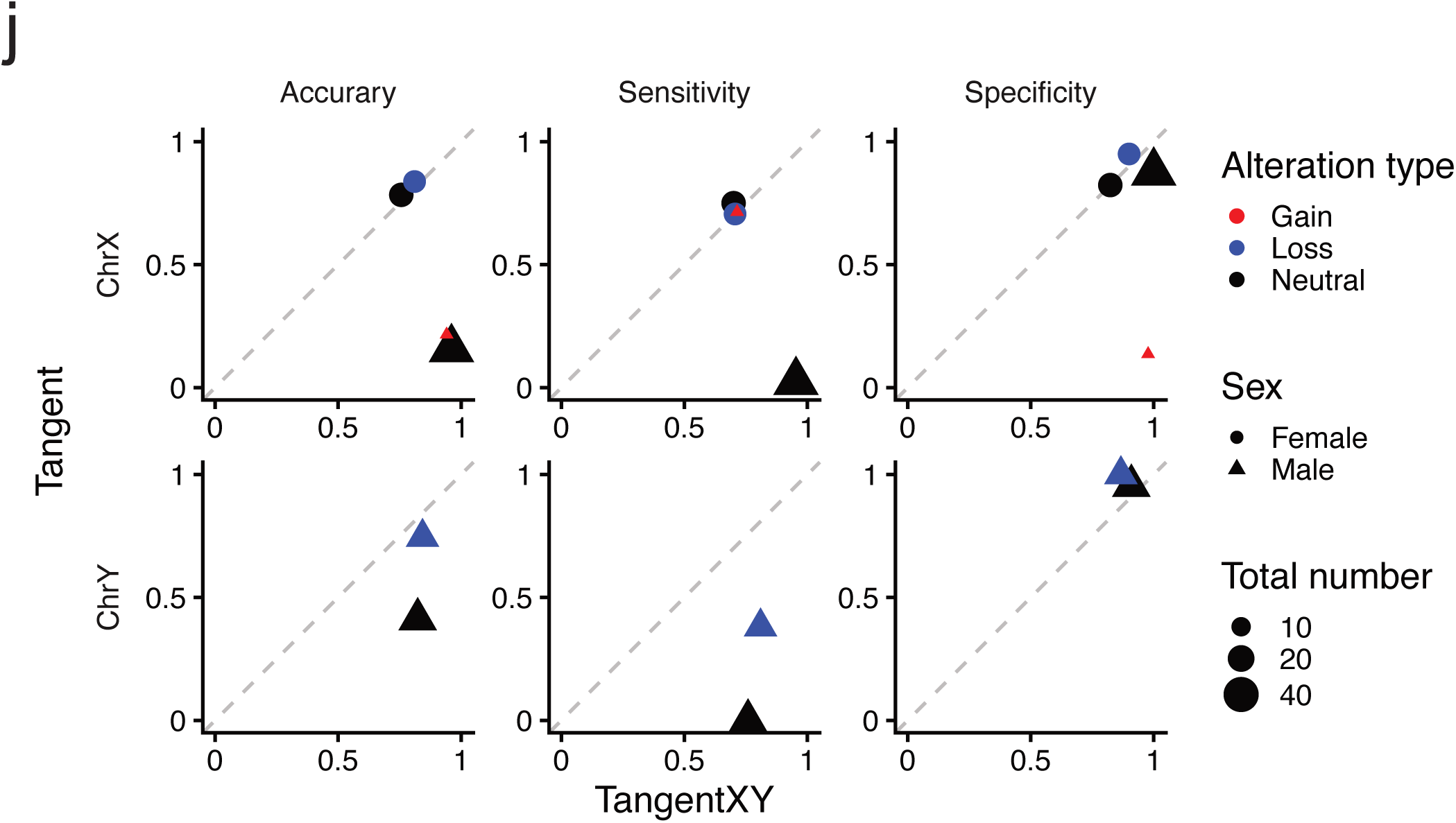
TangentXY and Tangent normalization on CCLE data. (**a-i**) Representative images of TangentXY/Tangent normalization on CCLE cell lines with validated karyotype annotations. The view is the same as Figure 2d **and 2e**. (**j**) Performance comparison between Tangent and TangentXY.

**Supplementary Figure 5.**
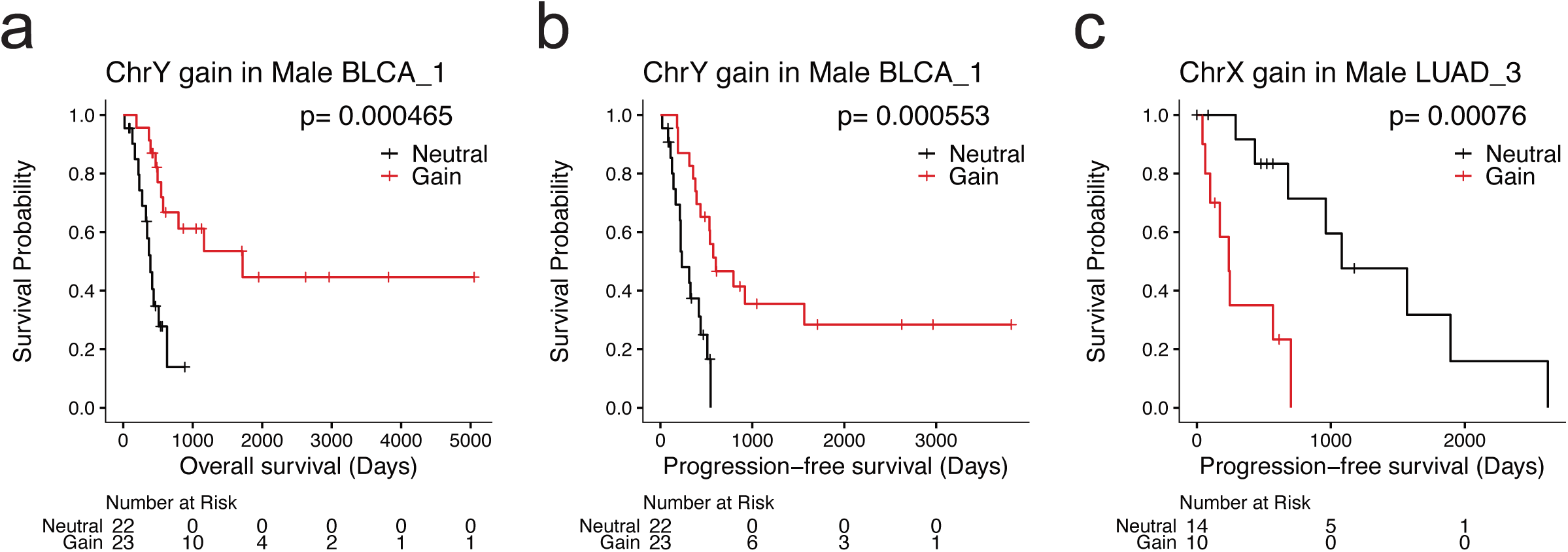
Kaplan-Meier plots of overall survival and progression-free survival across tumor subtypes. (**a**) OS in BLCA_1. (**b**) PFS in BLCA_1. (**c**) PFS in LUAD_3. P-values were determined by a log rank test.

**Supplementary Figure 6.**
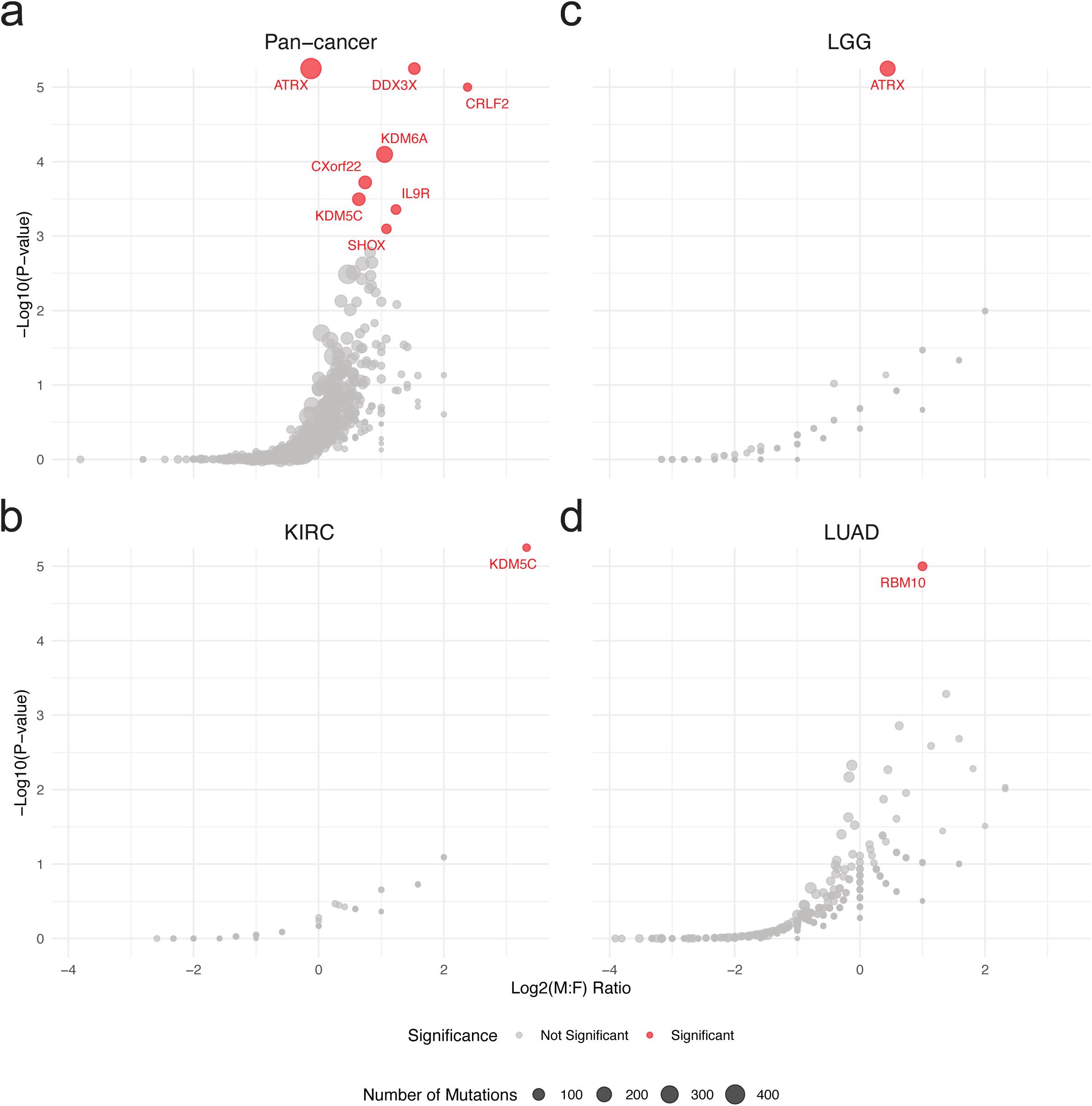
Male-to-female mutation ratios. Genes for which a statistically significant difference (adjusted p-value < 0. 05) in male-to-female mutation ratio was observed in the pan-cancer (a) and each cancer type (b-d) are shown in red.

**Supplementary Figure 7.**
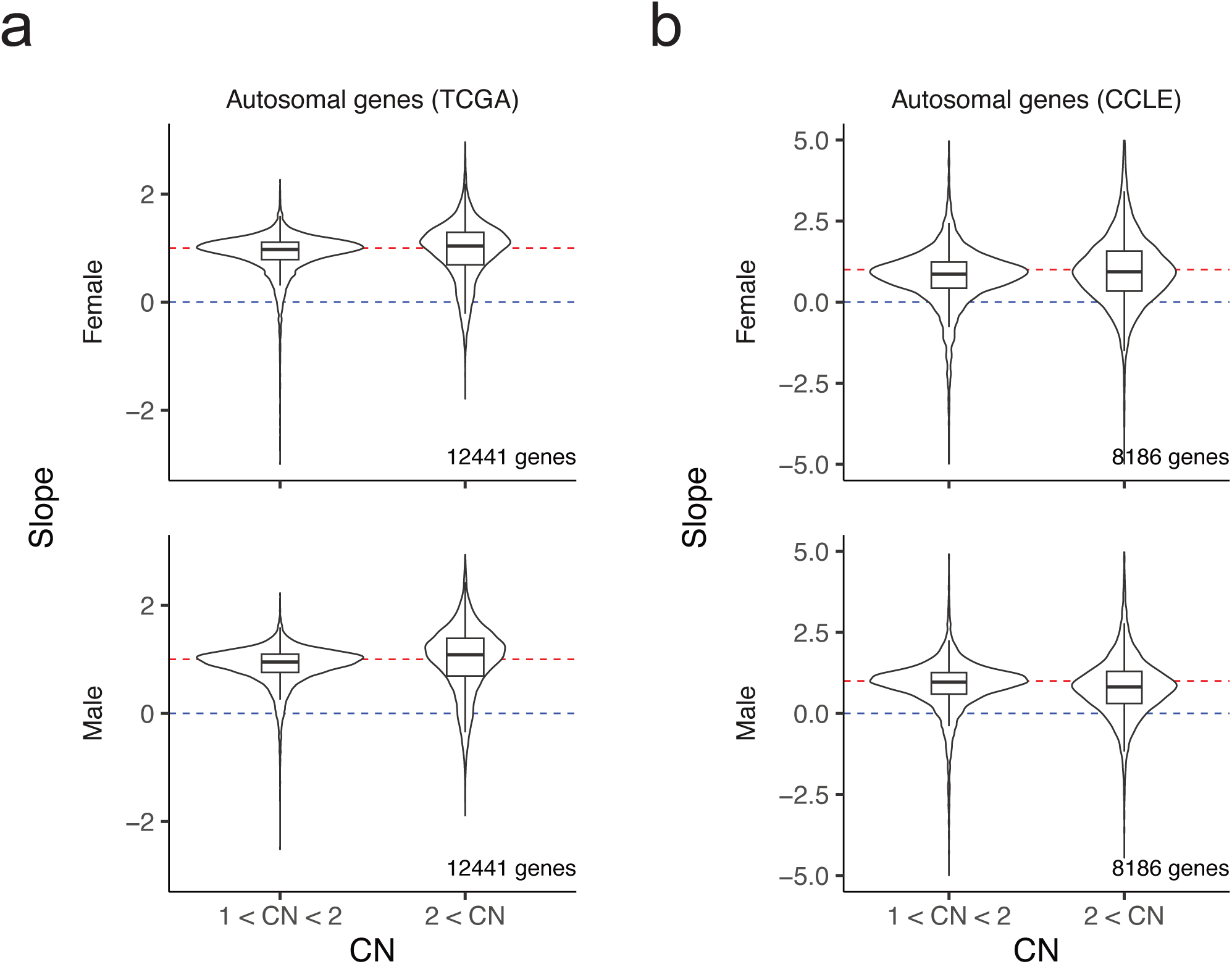
Slopes of autosomal genes. Slopes of autosomal genes in a region with copy-number less than 2 and greater than 2 in TCGA samples (**a**) and CCLE cell lines (**b**).

**Supplementary Figure 8.**
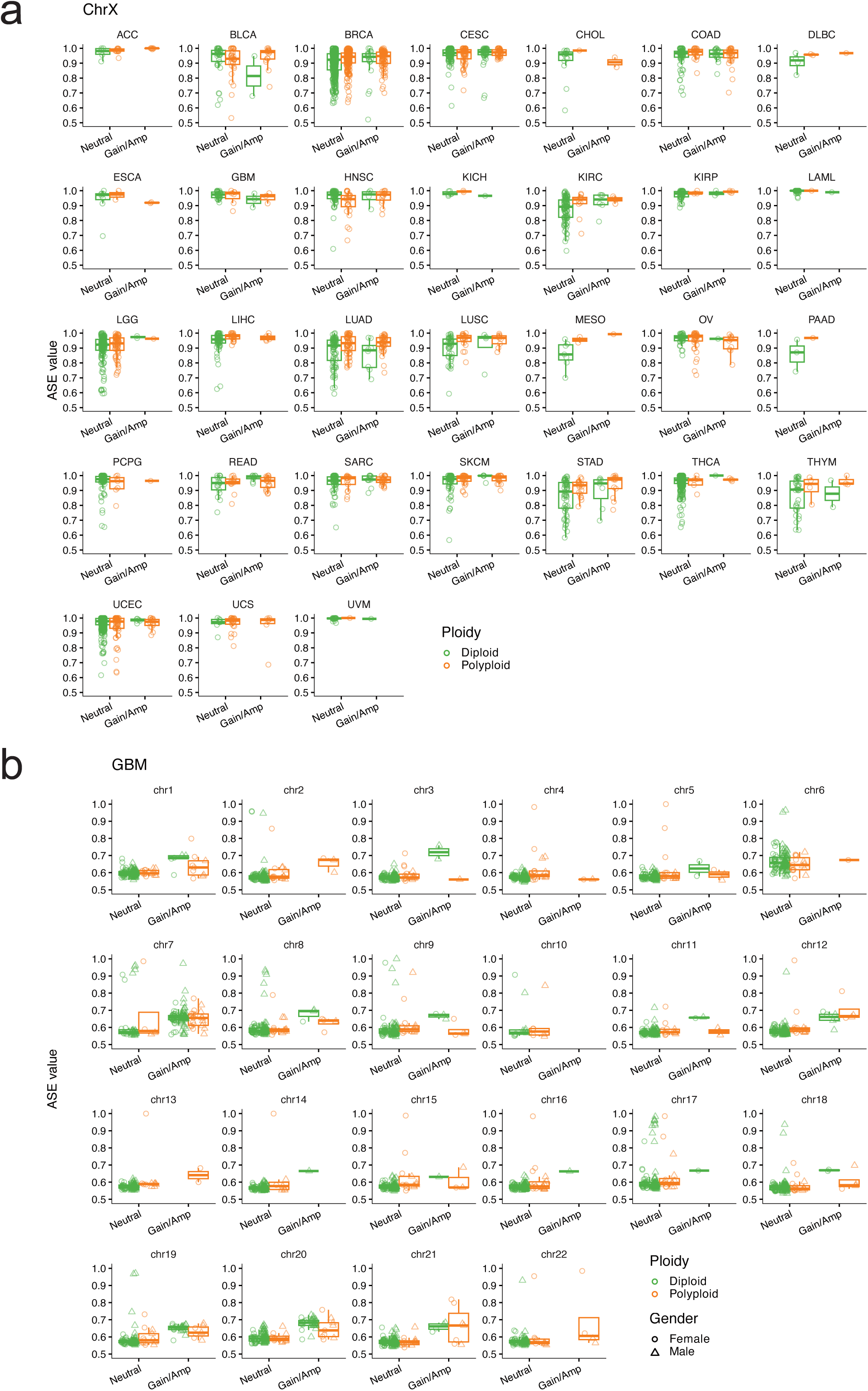
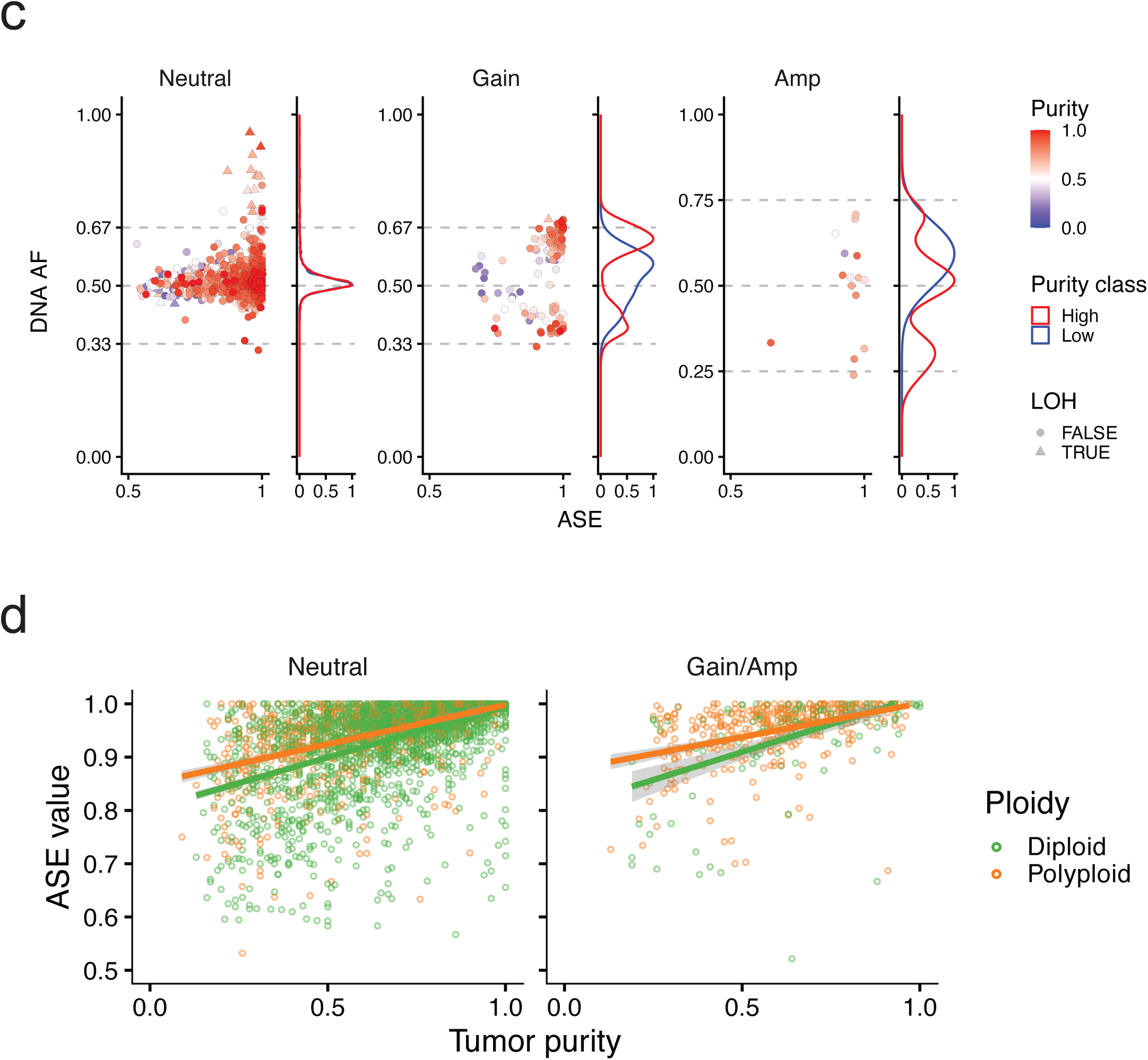
ASE of chrX in aneuploid cancer samples. (**a**) Median ASE values for TCGA samples shown in Figure 4e are separated by each tumor type. (**b**) Median ASE values for autosomal chromosomes in GBM. (**c**) Associations between DNA AF and ASE in diploid cells with different chrX SCNA status. (**d**) Associations between ASE value and tumor purity. The regression line is generated with a linear regression model.

